# Decoding semantics from natural speech using human intracranial EEG

**DOI:** 10.1101/2025.02.10.637051

**Authors:** Camille R. C. Pescatore, Haoyu Zhang, Alex E. Hadjinicolaou, Angelique C. Paulk, John D. Rolston, R. Mark Richardson, Ziv M. Williams, Jing Cai, Sydney S. Cash

## Abstract

Brain-computer interfaces (BCIs) hold promise for restoring natural language production capabilities in patients with speech impairments, potentially enabling smooth conversation that conveys meaningful information via synthesized words. While considerable progress has been made in decoding phonetic features of speech, our ability to extract lexical semantic information (i.e. the meaning of individual words) from neural activity remains largely unexplored. Moreover, most existing BCI research has relied on controlled experimental paradigms rather than natural conversation, limiting our understanding of semantic decoding in ecological contexts. Here, we investigated the feasibility of decoding lexical semantic information from stereo-electroencephalography (sEEG) recordings in 14 participants during spontaneous conversation. Using multivariate pattern analysis, we were able to decode word level semantic features during language production with an average accuracy of 21% across all participants compared to a chance level of 10%. This semantic decoding remained robust across different semantic representations while maintaining specificity to semantic features. Further, we identified a distributed left-lateralized network spanning precentral gyrus, pars triangularis, and middle temporal cortex, with low-frequency oscillations showing stronger contributions. Together, our results establish the feasibility of extracting word meanings from neural activity during natural speech production and demonstrate the potential for decoding semantic content from unconstrained speech.

## Introduction

Brain-Computer Interfaces (BCIs) represent a promising means for restoring communicative abilities in individuals with severe language production impairments (*1-6*). By circumventing the compromised neural-motor pathways, BCIs aim to directly translate the patient’s intended speech into meaningful output (*7, 8*). Over the past decade, the majority of BCI approaches have employed both invasive and non-invasive neural recording techniques to reconstruct various phonetic features of speech, including acoustic waveforms and phoneme sequences (*4, 9*). Such approaches have yielded increasingly fruitful results, achieving remarkable decoding accuracies in controlled experimental settings (*1, 7, 9-12*). In contrast, comparatively little research has focused on the reconstruction of semantics from neural signals. Semantic planning plays an important role, as the fundamental aspect of language production is to convey meaningful message between speakers (*13*), which allows for more robust and accurate communication even under challenging conditions like low signal quality or limited bandwidth (*14*). Semantic decoding, therefore, has the potential to provide natural, rapid and effective speech restoration.

Previous studies have demonstrated the feasibility of decoding semantic meanings from neural signals during both language comprehension and production using imaging methods, and have achieved notable accuracy in reconstructing the meaning of sentences (*15, 16*). However, these investigations primarily employed blocked experimental designs and lack the scenarios of natural language production. Natural language production during dialogue involves a diverse combination of words to convey a vast range of meanings, with temporal dynamics that can vary significantly based on the interactions between speakers (*17, 18*). As a result, decoding semantic features from natural language production in such contexts poses a considerably greater challenge (*19*). Due to these complexities, our understanding of how effectively lexical-semantic features can be decoded from language production in natural conversations remains limited.

Recent neuroimaging studies have revealed that semantic representation relies on a distributed network of interconnected brain regions (*20-25*). This semantic network includes regions in the temporal, parietal, frontal lobes, and subcortical structures, including activations in the anterior temporal lobe (ATL) and posterior middle temporal gyrus (pMTG) during various semantic tasks (*26, 27*). Further, the involvement of modality-specific sensory and motor areas suggests that semantic representations are partially grounded in the neural circuits that support language perception and production (*28, 29*). Together, this widespread, interconnected network underlies the complex nature of semantic processing in the human brain.

In this work, we employed a new approach by utilizing intracranial stereo-electroencephalography (sEEG) recordings during natural conversations to investigate the extent to which lexical semantic features can be decoded in language production (*30*). This approach allowed us to capture neural activity associated with spontaneous language production, providing a more naturalistic context. Using multiclass decoding of neural activity during natural speech production, we successfully predicted lexical semantic features across all participants, demonstrating the feasibility of reliably decoding lexical semantic information from neural activity during natural language production. Key brain regions and oscillations in specific frequency bands were also identified. Together, our findings pave the way for future research into the neural basis of semantic processing in real-world communication contexts.

## Results

### Neural recording during natural language production

In this work, we utilized semi-chronically implanted depth sEEG recordings from 14 participants undergoing epilepsy monitoring as part of their clinical care (6 females and 8 males, average age of 34, ranging between 16 to 59, **Figure 1A**) (*30*). Recording channels were distributed across the frontotemporal lobes, covering major brain areas known for language production planning, including superior and middle temporal cortices, precentral and pars triangularis (**Figure 1B**). Voltages recorded from these channels were filtered to six frequency bands, including theta (4-8 Hz), alpha (8-12 Hz), beta (12-30 Hz), low-gamma (30-55 Hz), mid-gamma (70-110 Hz) and high-gamma (130-170 Hz), and the envelopes of each band were calculated.

**Figure 1.**
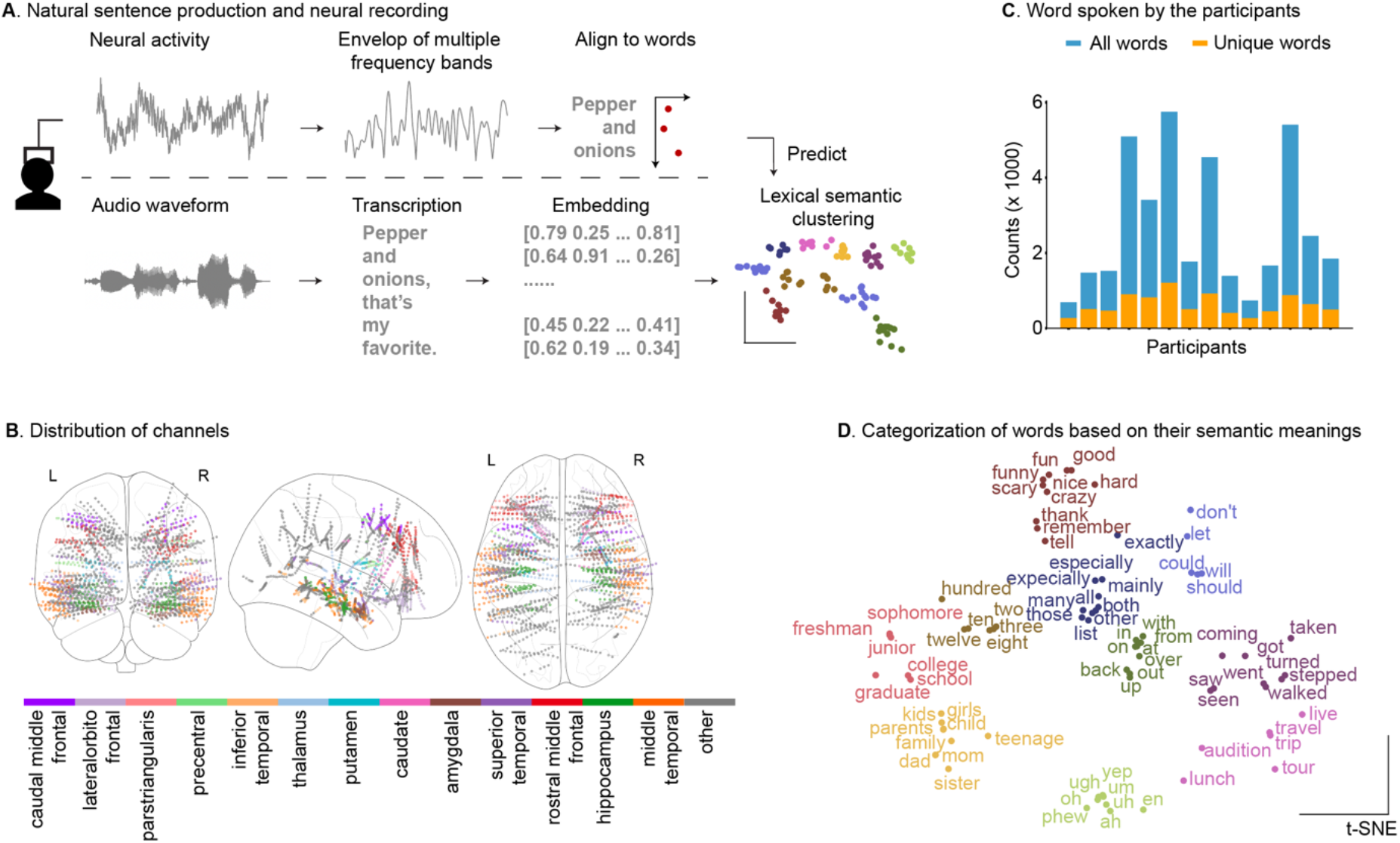
Natural speech production and channel distribution. **A**. Participants engaged in natural conversations while their neural activity was recorded by stereo-electroencephalography. Neural activity was bipolar-referenced and filtered to various frequency bands, and the envelope of each band was extracted. Meanwhile, the words the participants produced were transcribed and temporally aligned with the neural recordings. The lexical semantic content of participants’ speech was encoded using word2vec embeddings and subsequently categorized into 10 distinct semantic clusters to capture different word meanings. **B**. The recording included a total of 1908 channels, which were primarily distributed in the frontal, temporal and mesial areas in both hemispheres. **C**. The total number of words and unique words are plotted by each participant. On average, each participant produced 2,696 words, with 621 unique words among them. **D**. To categorize the meaning of the words, we performed spectral clustering on the word2vec embeddings into 10 categories. To visualize this semantic organization, we projected selected words from one participant onto their first two t-SNE dimensions, illustrating the natural clustering of meaning in this reduced dimensional space.

To allow for the most ecological speech production, participants engaged in natural conversations with an experimenter, which lasted between 16 to 92 minutes. From these conversations, we transcribed words in sentences during speech production. In total, each participant spoke an average of 2,696 ± 1,786 (Mean ± s.t.d) words, including 621 ± 278 unique, unrepeated words (**Figure 1C**). The sentences cover a wide range of topics, such as personal experience, movies, video games, career, family and travel plans (**Table S1**).

### Lexical semantic categorization

In natural conversation, lexical semantic meaning can be highly diverse, so we first represented these semantic features using vectorized representations to enable more accurate decoding of neural activity. Word2vec represents a powerful approach for encoding lexical semantics through dense vector representations (*31*). By learning from extensive text corpora, word2vec captures semantic relationships between words based on their contextual co-occurrence patterns. This methodology enables words with similar semantic content to cluster together in a high-dimensional vector space. The ability to encode lexical semantic information in a vectorized format makes word2vec an ideal tool for examining whether neural activity reflects underlying semantic structures. In our analysis, we extracted 300-dimensional word2vec embeddings for each word uttered by participants, and excluded words not present in the pre-trained vocabulary.

Given the open-ended nature of our dataset, which allowed participants to freely produce language without lexical constraints, comprehensively characterizing the semantic content of the thousands of recorded words presents a significant analytical challenge. To mitigate this complexity, we implemented a dimensionality reduction approach by categorizing words into 10 semantic clusters using word2vec embedding representations. This number of clusters ensures that each category captures a meaningful portion of the vocabulary, with most containing between 2% and 20% of the produced words. The resulting clusters were comprised of words with similar meanings. For example, ‘sisters and ‘family’ were in the same cluster, whereas ‘scary’ and ‘travel’ were in separate clusters (**Figure 1D**). In addition, this clustering procedure produced generally stable results across multiple runs. On average, 68.8% of the words were consistently grouped into the same clusters across different clustering iterations, regardless of the initial random seed used to start the clustering process. This high level of consistency suggests that the clustering method is reliable. The exact words that were consistent across different iterations were shown in **Table S2**. In this way, we categorized words reliably based on their lexical semantic representation.

### Lexical semantic decoding from local field potentials

To investigate to what extent the semantic category of individual words can be readout from neural activity during natural sentence production, we developed a multivariate decoding model to predict the identity of 10 clusters of lexical semantics. This approach allowed us to test whether patterns of neural activity contained sufficient information to predict the semantic feature of each word. Since we had 10 possible clusters, chance performance would be 10% if the model made predictions randomly. However, using a 5-fold cross-validation framework where the model was trained on 80% of words and tested on held-out words, we found that our decoding model reached a decoding accuracy of 20.9 ± 3.4% (mean ± s.t.d) across 14 participants (**Figure 2A**). Notably, some participants achieved a decoding accuracy as high as 26.50%, which was 2.65 times higher than what can be explained by chance. The decoding accuracy across different semantic clusters also displayed variations (Standard deviation across clusters ranges from 7% to 19%, with a mean of 13%, **Figure S1**). Next, we compared the performance against a permutation control where we randomly shuffled the word labels relative to the neural data, thereby destroying any true relationship between neural patterns and word meanings while preserving the distribution of words across clusters. This permutation analysis yielded an average accuracy of 10.0%, confirming that our observed decoding performance significantly exceeded chance levels (p = 0.0001 for all participants) and could not be explained by any imbalance in the number of words per semantic cluster.

**Figure 2.**
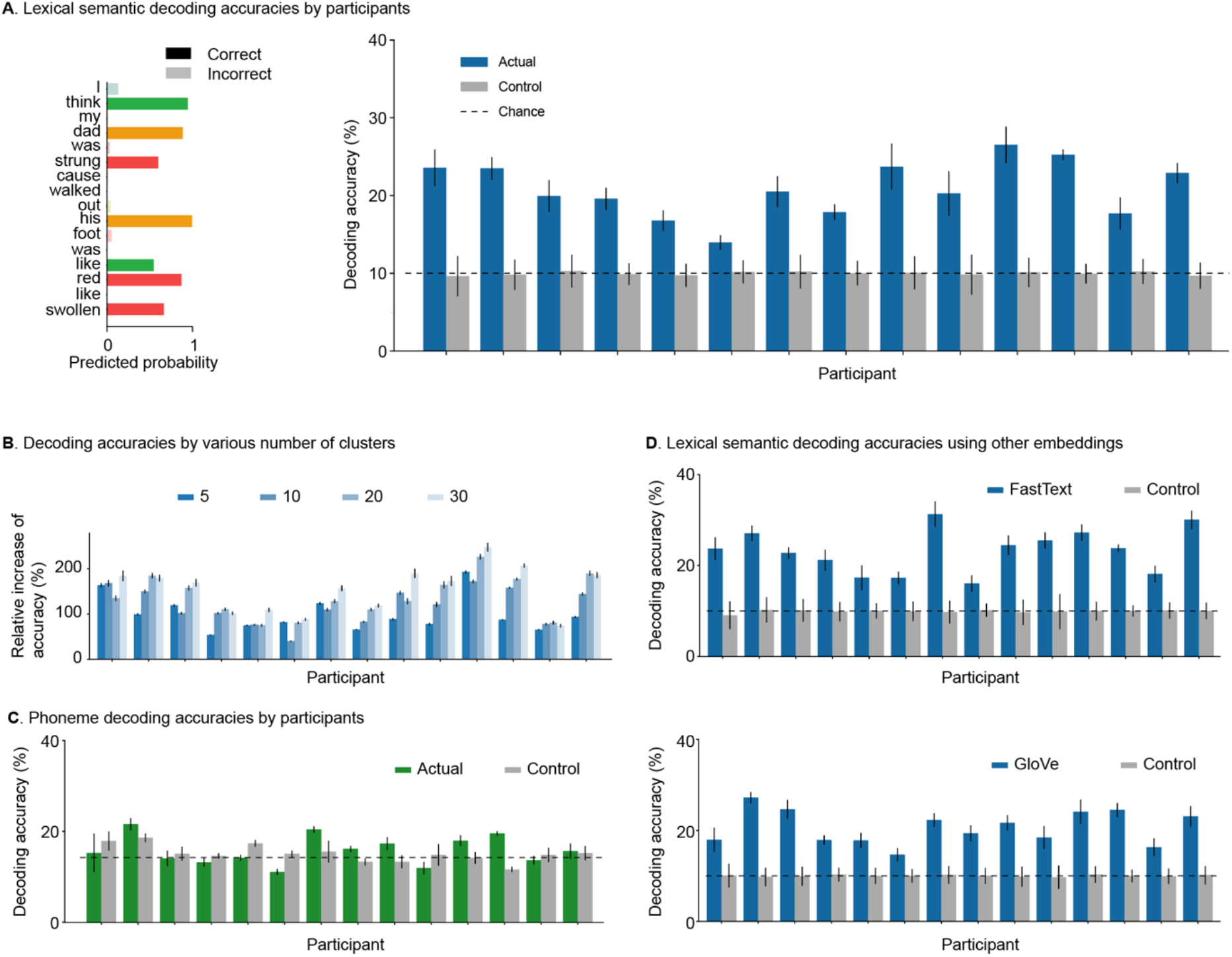
Lexical semantic decoding results and control analysis. **A**. *Left*, example sentence with decoding results for each word. The color labels the cluster of the word, whereas the transparency labels whether the word was predicted correctly or not. *Right*, single-participant decoding accuracies (blue) are shown with the performance derived from randomized permutation controls (gray). The theoretical chance level was determined by the reciprocal of the total number of clusters. **B**. The percentage improvement in decoding accuracy compared to chance remained consistent across different numbers of clusters. **C**. Decoding accuracy for phoneme clusters remained at chance level, indicating that phonetic planning features did not drive the result in lexical semantic decoding. **D**. Comparable decoding accuracies were achieved using fastText (top) and GloVe (bottom) embeddings.

Next, to ensure that the decoding capability of the neural activity was not limited only to the specific number of lexical semantic clusters, we performed the same modeling procedures, but using various number of word clusters of 5, 10, 20 and 30 clusters (**Figure 2B, Figure S2**). Overall, the decoding accuracies were significantly higher than chance for all the numbers of cluster we tested (T-test, p ≤ 1 × 10^−7^). Further, despite a decrease of absolute decoding accuracy with the increase of cluster number, the relative increase of accuracy compared to chance remains consistently higher than chance among 5, 10, 20 and 30 clusters, ranging from 99.1% to 156.0%. This suggests that the decoding feasibility is robust across varying levels of granularity in the semantic clustering.

We further investigated the temporal consistency of lexical semantic decoding by examining whether the decoding remained stable across different neural activity time windows. Utilizing the same decoding methods, we analyzed neural activity averaged across window durations ranging from 100 to 900 milliseconds prior to word onset. The average decoding accuracy across participants consistently ranged from 19% to 23%, with no statistically significant differences between temporal windows (One-way ANOVA, p = 0.21, **Figure S3**). These findings suggest that lexical semantic neural representations are not confined to a specific temporal duration but rather exhibit a consistent pattern of neural encoding across multiple time intervals.

To rule out the possibility that lexical semantic decoding results arise from neural activity related to the motor planning of word pronunciation, we conducted a control analysis to decode the phonetic features of each word. Specifically, we applied identical decoding procedures to seven consonant clusters for each word. We found, however, that the decoding accuracy for phonetic features (15.9% ± 3.0%) was statistically indistinguishable from chance levels (15.1% ± 1.8%, Permutation test, p = 0.22). These results demonstrate that our semantic decoding is not driven by neural representations of word pronunciation, thereby suggesting the specificity of semantic feature encoding in neural activity.

Last, we examined whether the decoding capability was generalizable across different lexical semantic embeddings. Instead of using word2vec, we performed the decoding analysis on categories based on two independent word embeddings, FastText and GloVe, and observed similar decoding accuracies from these two embeddings (FastText: 23.3% ± 4.6%; GloVe: 20.7% ± 3.6%), with all participants showing significantly higher than chance decoding (Permutation test, p = 1 × 10^−5^). Further, the decoding accuracies were similar across these three embeddings (ANOVA, p = 0.18), indicating that the prediction of lexical semantic was not dependent on the specific word embeddings.

### Lexical semantic computation across various brain areas and frequencies

Next, we identified the specific brain regions and frequency bands contributing to the decoding of lexical semantic clusters to investigate the specific neural basis of the lexical semantic decoding. Based on the selection criteria of statistically significant responses to lexical semantic features using the Wilcoxon rank-sum test (p-value < 0.05, Bonferroni corrected for clusters), we found an average of 12.1% channels showing selective responses across participants (**Figure 3A**). This selectivity was significantly higher than those from chance in all participants (Chi square proportion test, p < 0.05), with majority of channels and frequencies displayed selective responses to only one semantic category (91% of responding channels, **Figure S4**). Together, these findings suggest that lexical semantic information is encoded through the coordinated activity of these selectively responding channels.

**Figure 3.**
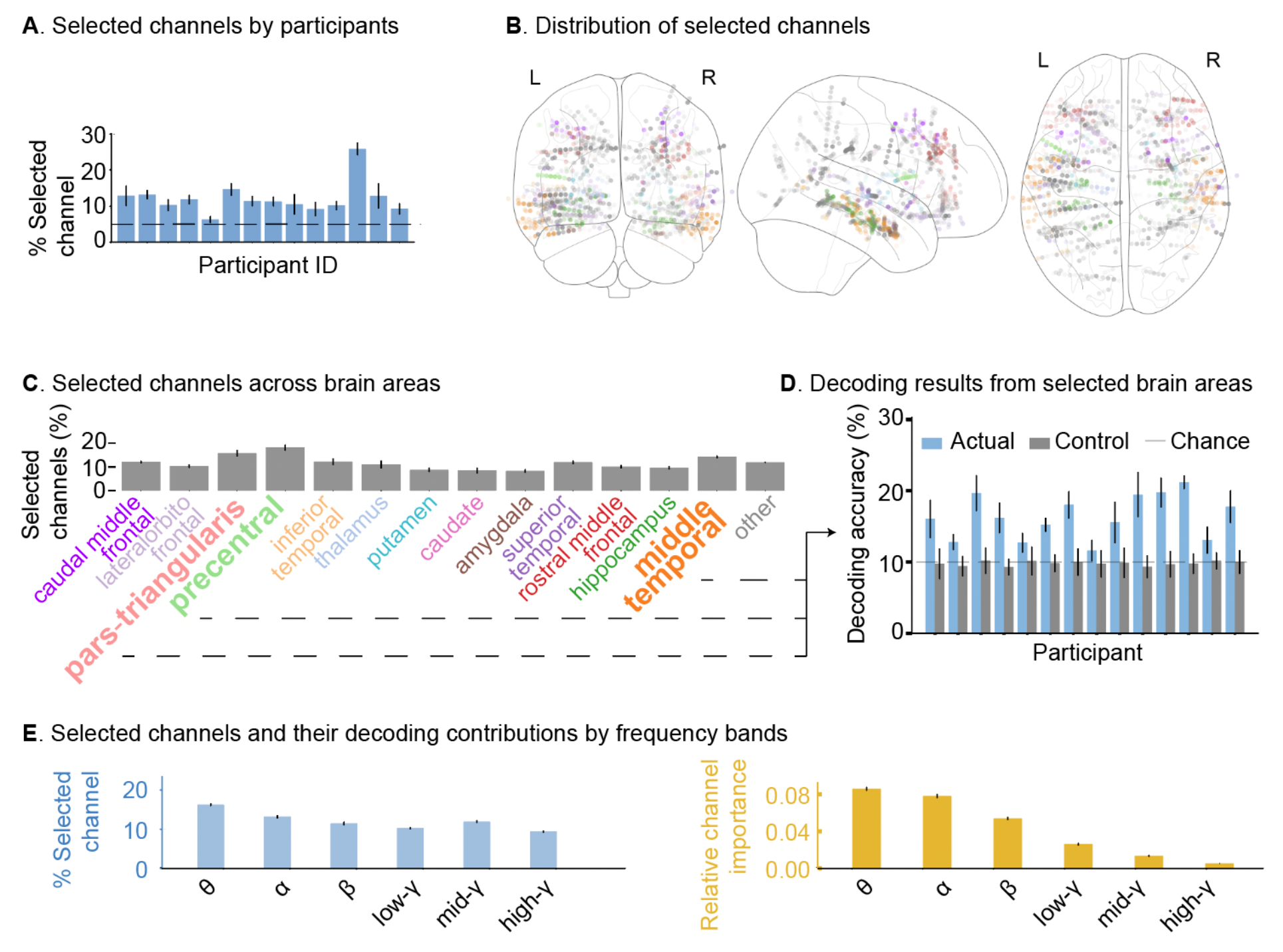
Neural contributions to lexical semantic decoding across brain regions and frequency bands. **A**. Percentage of selected channels across participants. **B**. Distribution of selected channels across various brain areas. The transparency of each channel indicates how frequently this channel is selected across different clustering iterations. **C**. The percentage of selected channels across various brain areas. Among these areas, we identified that pars-triangularis, precentral and middle temporal cortex (bolded) showed significantly higher percentage of responding channels than other areas. **D**. The decoding accuracy using channels from these three areas. **E**. Distribution of neural signal contributions across frequency bands in the selected channels. Blue bars represent the percentage of channels within each frequency band, while orange bars indicate the relative importance of channels based on their feature importance in each frequency range.

The channels that responded to lexical semantic were distributed broadly across multiple brain areas (**Figure 3B, C**). Not surprisingly, we found that the left hemisphere was more involved in semantic planning compared to the right hemisphere (T-test, p = 1.63x 10^−7^). This lateralization effect is, as expected, consistent with expectations of language lateralization (*32*). Further, we focused on identifying the brain areas with a high number of recording channels across participants (> 30 channels), which resulted in 13 specific brain areas. To determine which of these high-coverage areas were significantly different from the rest, we performed a T-test comparing each area to the others. The result revealed that middle temporal cortex, precentral and pars triangularis showed significantly higher ratio of responding channels than the other brain areas (T-test, p < 0.05, **Figure S5**). Together, by examining neural response during language production planning, we found a specialized yet distributed nature of semantic representation in various brain areas.

Based on the identification of semantic processing brain areas, we next investigated the extent to which lexical semantic features could be decoded using only these targeted regions. Decoding analysis using channels only from these three regions showed an accuracy of 16.4 ± 3.6% (mean ± s.t.d), which were significantly higher than those predicted by chance (Permutation test, p < 10^−5^, **Figure 3D**). However, decoding accuracy from channels in these three areas were significantly lower compared to decoding accuracy using channels from all areas (Permutation test, p = 0.006). Together, we found that the semantic processing areas contributed significantly to lexical semantic decoding, though including broader area will increase the decoding accuracy.

To further understand the neural mechanisms underlying lexical semantic decoding, we analyzed the distribution of neural responses across different frequency bands. When examining the selected neural responses, we found that all investigated frequency bands (theta, alpha, beta and gamma bands) were present but that there was greater representation in lower frequencies (ANOVA, p = 9 × 10^−78^, **Figure 3E**). A similar trend was also observed when considering the feature importance of different frequency bands, revealing a strong negative correlation (**Figure 3E**, Pearson correlation coefficient, R = -0.83, p = 0.043). This low-frequency bias was not an artifact of our time window selection method, as using smaller windows yielded consistent results (**Figure S6**). Together, these findings collectively illustrate how a diverse population of neural channels and specific frequency bands collaboratively contribute to the planning of lexical semantic production, suggesting that lower frequency neural oscillations play a particularly significant role in semantic information processing.

## Discussion

While recent advances in neural technology and computational methods have enabled the decoding of various aspects of language from neural signals (*1-10*), the ability to extract word-specific semantic content during natural speech production has remained elusive. Here, we demonstrate the successful decoding of lexical semantic information from sEEG recordings in 14 participants engaged in spontaneous language production during natural conversation (*30*). Our analyses revealed consistent above-chance decoding accuracy across participants, with robust performance across multiple semantic clustering and embedding models. Importantly, control analyses showed chance-level decoding of phonemic features, indicating that little phonetic features were encoded in these channels, therefore the semantic decoding results was not driven by these low-level acoustic properties. These findings provide compelling evidence for the neural encoding of lexical semantics while producing natural speech and establish the feasibility of extracting word meanings from human brain activity during unconstrained language production.

Natural conversation represents a uniquely challenging context for studying language production mechanisms, as it involves potentially richer semantics and temporal dynamics compared to controlled experimental paradigms (*17, 18*). During spontaneous discourse, speakers freely select topics, vocabulary, and semantic content while engaging in dynamic linguistic behaviors such as co-construction and backchanneling (*33*). These naturalistic elements introduce substantial variability that has traditionally complicated the systematic study of language processes. Nevertheless, our striking ability to achieve above-chance decoding accuracy of lexical semantic features under these unconstrained conditions demonstrates the robustness of semantic representations in neural activity during natural communicative interactions. While this level of decoding is insufficient for any clinically useful decoding system, this success in decoding semantic content from natural speech extends previous findings from more controlled experimental settings to ecologically valid contexts (*16*). It also suggests that more intentionally diverse and complete cortical coverage may permit even greater decoding accuracy.

Another striking finding is that we showed the neural substrates supporting lexical semantic processing during natural language production comprise an extensive, interconnected network with left hemisphere lateralization (*21, 32*). Across this distributed system, we identified regions that contributed disproportionately higher to semantic decoding including the precentral gyrus, pars triangularis, and middle temporal cortex. This suggests that semantic representations during speech production are predominantly, although not exclusively, encoded by areas known to process language and motor planning (*34-38*). However, we showed that semantic planning extends beyond these language areas: The decoding accuracy from all regions was significantly higher than those from the three major regions, suggesting that semantic information is distributed across multiple cortical regions rather than being confined to traditional language areas. The distributed nature of semantic processing suggests that optimal semantic BCI performance may need to incorporate information from a broader network of brain areas to capture intended meaning during speech production.

Further, these results indicate that semantic information was encoded across a broad frequency range, from theta to gamma bands. Notably, lower frequencies yielded higher decoding accuracy gain compared to higher frequencies, potentially reflecting semantic retrieval processes (*39-41*). This preference to lower frequencies was not accounted for by the time window used to calculate neural activity, as smaller time windows resulted in similar trends. Although the type and design of recording electrodes may impact the contribution of frequency bands (*42*), these findings indicate that lexical semantic processing during natural speech production is supported by a distributed neural network across multiple frequency bands, with particularly strong contributions from lower-frequency oscillations.

Together, our findings establish the feasibility of decoding lexical semantic information during natural language production and provide a detailed mapping of the neural substrates supporting this process. However, there are several limitations for this study: First, our study prioritized demonstrating the basic feasibility of semantic decoding over maximizing word-level accuracy. Future investigations employing larger word samples per participant may achieve higher decoding precision and enable more fine-grained semantic categorization. Further, we observed substantial inter-individual variability in decoding performance, and the contributions of electrode location variation versus genuine individual neural coding differences remain to be revealed.

Despite these limitations, this work represents a crucial first step in decoding lexical semantic features during natural speech production and establishes a foundation for future research in this domain. Most importantly, we envision that this approach may be leveraged with other models of speech decoding, particularly those which are phoneme focused, to produce ever smoother, more accurate and rapid abilities to restore communication to those who have lost it due to severe neurological injury.

## Acknowledgement

C.R.C.P is supported by Madame Stella Zorzet and the Zorzet legacy through the EPFL WISH Foundation. Z.M.W. is supported by NIH R01DC019653 and NIH U01NS123130. J.C is supported by the Mussallem Transformative Award and American Association of University Women. S.S.C. is supported by NIH U01NS098968. We utilized Anthropic Claude to refine the language of this manuscript.

