## Supplementary Material for "Decoding semantics from natural speech using human intracranial EEG"

### Materials and Methods

#### Stereo-Electroencephalography recording

In this study, we leverage intracranial stereo-electroencephalography (sEEG) recordings from 14 participants recorded in our previous study (*1*). Recordings of neural activity and speech were obtained with informed consent from patients with intractable epilepsy (6 female, 8 male, 10 right-handed, age from 16 to 59) undergoing seizure localization (Massachusetts General Brigham Institutional Review Board). All participants had normal speech and comprehension ability. Before enrolling, participants were fully informed that their involvement in the study would not affect their clinical care, and that they could withdraw at any time without any impact on their treatment. The procedure for implanting electrodes was carried out independently of the study itself, ensuring no conflict of interest. Informed consent was obtained from all participants prior to their inclusion in the study.

Neuronal activity (128-256 channels, NSP/Cerebus/Blackrock) was recorded at 2 kHz and filtered online (1 kHz) for local field potentials. Neural data was re-referenced to bipolar configuration using MATLAB (MathWorks) and NPMK toolbox (Blackrock). No seizures occurred during conversation, but interictal epileptiform discharges (IEDs) were identified using in-house MATLAB code based on Janca et al *(2)* and channels with high rate of IED were excluded, resulting in 1908 bipolar channels for analysis. Next, the envelope from Hilbert transformation was extracted for theta (4-8 Hz), alpha (8-12 Hz), beta (12-30 Hz), low-gamma (30-55 Hz), mid-gamma (70-110 Hz), and high-gamma (130-170 Hz) bands (*1*). Neural activity was further processed to remove outliers and linear detrended to remove neural drifts over recording (*1*).

#### Natural speech production and audio recording

During neural recordings, participants were engaged in a free conversation with an experimenter (**Figure 1A**) (*1*). During the conversation, participants produced an average of 2,696 ± 1,786 (mean ± s.t.d) words (excluding non-meaningful words such as 'ha'). Among these words, there were 621 ± 278 unique, unrepeated words. Topics ranged from informal discussions (favorite food, travel plans) to events (daily schedule, health status) between experimenter and participant (**Table S1**). Audio was recorded on a DR-40X recorder (TASCAM) and synchronized to neural data via the NSP system (Blackrock). Audio was firstly transcribed with WhisperX (*3*), then manually corrected using Audacity (v3) or SpeechScribe (in-house Python tool). Participants' personal information (names, ages, etc) were replaced by standardized identifiers to ensure participants privacy.

Following speech transcription, words were temporally aligned to the corresponding neural activity. For each word, we averaged the preceding 500 milliseconds of the neural envelope signal (extracted from six frequency bands per channel) to create a characteristic neural activity representation. This window of time balances the capture of neural processing of speech production planning while minimizing overlap with preceding words.

#### Lexical semantic representation and classification

Word2vec is an effective method for representing the lexical semantics of words through dense vector representations (*4*). In our analysis, we used the word2vec model that was pretrained on a 100 billion word-subset of the Google news dataset. Specifically, we obtained the 300-dimension word2vec embeddings from each word that the participant uttered, and removed words that were not present in the word2vec pre-trained vocabulary.

Given the limited number of words produced by each participant, it is unlikely to predict the exact lexical semantics for each specific word. Instead, we categorized words into clusters based on their lexical semantics and performed decoding models to predict the word-meaning category. Specifically, we clustered words based on their word2vec embeddings using spectral clustering algorithm (*5, 6*). Spectral clustering was chosen because it is a hypothesis-free and efficient method for high-dimensional data, allowing for embeddings that are not normally distributed or with different covariance matrices among clusters (*7*). Spectral clustering works by transforming the original data into a lower-dimensional space where clusters are more easily separable. This transformation is based on the spectrum (eigenvalues) of the Laplacian matrix of the similarity graph between instances:

$$\text{similarity}(x, y) = e^{\frac{-||x-y||^2}{2\sigma^2}}$$

with $||\cdot||$ being the Euclidean distance and $\sigma$ the parameter controlling the width of the kernel. In this work, spectral clustering is performed on the z-score normalized word2vec embeddings from a subset of unique words from the vocabulary produced by participant. Further, a cluster number of 10 was fixed for each patient. This number was chosen with the consideration of having a maximum cluster number in the constraint of having few clusters with only few words in them. We manually verified that the proposed clustering ensured cluster interpretability, i.e. clusters were grouping semantically similar words (**Figure 1D**). The clustering procedure was repeated for 20 different fixed seeds to ensure both stability and reproducibility of the results.

To assess the generalizability of our findings beyond a single embedding model, we included two additional semantic word embeddings models: fastText (*8*) and GloVe (*9*). FastText, an extension of word2vec, incorporates morphological information to better capture word complexity, while GloVe, a log-linear model, focuses on co-occurrence ratios to maximize word-pair similarity and mitigate noise. These two pre-trained models were also trained on different datasets. Finally, we compared decoding performance across all the word embeddings using a one-way ANOVA test to identify statistically significant differences.

#### Selecting channels that show significant response to lexical semantics

To identify neural activity most relevant to semantic processing during speech production, we implemented a feature selection procedure to obtain a smaller number of neural dimensions. From the neural activity containing six frequency bands from all channels, we independently selected channels and frequency bands that displayed statistically significant selectivity for the semantic clusters using Wilcoxon ranksum one-vs-all tests (*10*). A threshold of 0.05 was used with Bonferroni correction for multiple clusters to decide whether a channel / band shows significant response to lexical semantics. The channels and the frequency bands were independently examined without introducing prior assumptions of the importance of specific frequency bands in speech production. To ensure the robustness of our selection results, we repeated the categorization process 20 times, each iteration using a different random seed.

To assess whether the time window length influenced a channel's frequency band response to lexical semantic features, we applied the selection method across multiple time windows spanning 5 to 500 milliseconds. For this analysis, we utilized a single random seed.

#### Semantic decoding model

To evaluate to what degree lexical semantic clusters can be deciphered from the neural activity, we employed multinomial logistic regression for speech decoding. Specifically, we used channels and frequency bands that showed significant response and predicted the cluster of the lexical semantic for each word. Each frequency band from each channel was considered as a single independent feature (*11*), and a limited memory Broyden-Fletcher-Goldfarb-Shanno (LBFGS) solver with an L2 regularization was used. For each participant, observations were first split into training and testing sets using a stratified 5-fold split. The decoding procedure was repeated for semantic clusters generated for different clustering seeds. Then we used bootstrapping method to resample training and testing data independently to account for potential noise in the neural recordings. Decoding accuracy due to the imbalance of cluster and randomness from neural activity was estimated by randomly permuting the clusters and repeating the above decoding procedures.

To examine how the granularity of the lexical semantic clustering impacts decoding results, we performed the decoding analysis with varying numbers of clusters. Specifically, we extended the number of clusters from 10 to 5, 20, and 30, and calculated the relative increase in decoding accuracy compared to random permutation for each participant:

$$\text{Relative Decoding Accuracy} = \frac{\text{Actual - Shuffle}}{\text{Shuffle}} \cdot 100\%$$

To evaluate the potential influence of pronunciation of words on decoding accuracy, we conducted a control analysis using the neural activity to predict the word phonemes (*12*). Specifically, we obtained the phoneme categories of consonants (Plosive, Fricative, Affricate, Nasal, Lateral, Rhotic, Glide) for the first phoneme of each word. We repeated the same procedures described above to calculate the decoding accuracy using neural activity.

In addition, to evaluate the contribution of channel and frequency band to the decoding accuracy, we calculated feature importances for the independent variable by estimating the relative increase of the accuracy of the variable: We first calculated the decoding accuracy while a specific independent variable was replaced by a randomly permutated one, and then we divided this value by the decoding accuracy obtained including all independent variables. We used the relative increase of the decoding accuracy with the consideration of variations of decoding results across individuals. Next, we selected the maximum feature importance across all iterations and standardized the feature importance score by dividing the decoding accuracy.

**Table S1. Example of sentences produced by the participants**

| |
|---|
| It’s an old town and nothing really gets updated. |
| Yep, first time out of the country so I can’t wait. |
| I got my engineering degree from here then I went back home. |
| I love process development. I think it’s awesome. |
| I have zero brother and zero sisters. |
| I did that because I was in my first love of my life, and she told me get a job in another job and then she told me quit the job. |
| It was truthfully scary. It was like watching a movie. |
| I’d also go out and travel like what you said visit, uh, the different countries that I want to see. |
| There was a Lego movie before, but not a Batman Lego movie. |

**Table S2. Example of categorization of words based on their lexical semantic meaning**

| |
|---|
| child, dad, family, girls, kids, mom, parents, sister, sisters, teenage, phone, them, people, someone, singer, die, my, she, hours, him, age, younger, doctor, young, animals, her, me |
| audition, live, show, shows, tour, tours, travel, trip, welcome, experiences, jazz, choir, concert, hum, lunch, dinners, eat, chorus, choirs, swim, courses, offers, watch, classes, attracts, fill, interested |
| mouse, fish, sharks, ocean, lake, field, water, sea, pool, day, swollen, red, foot, his, strung, apart, intense, she’s, literally, same, per, bizarre, closer, first, sca, super, urchin, between, regular, where, little, assumes, third, around, tiny, hand, late, head, on, out, just, right, before, body, top, middle, half, hotel, stick, rest, next, turns, when, getting, at, back, room, high, big, every, then, through, moving, makes, an, basically, into, once, seal, end, spot, there’s, slightly, each, from, in, over, up, with, almost |
| ten, four, sixteen, three, five, six, two, eight, hundred, twelve, thousand, seven, eighteen, twenty, fifteen, eleven, fourteen, few |
| crazy, fun, feel, funny, good, hard, nice, hate, love, scary, sorry, tell, thank, honestly, maybe, sense, looking, kinda, mind, done, hey, play, know, guess, hear, bunch, clue, knows, loved, bit, fortunate, seeing, mickey, watching, different, always, kind, memories, remember, look, probably, thing, something, here, really, very, there, yeah, see, think, personality, like, stuff, lot, everywhere |
| college, colleges, collegiate, freshman, freshmen, graduate, graduating, junior, school, sophomore, recruiting, coaches, team, seniors, players, coach, lacrosse, recruit, academic, academics |
| coming, got, saw, seen, stepped, taken, traveled, turned, walked, went, thought, has, been, was, knew, weeks, years, wrestled, months, summer, missing, said, quit, had, since, stung, called, say, exaggerating, snorkeled, year, last, days, bothered, minutes, lived, fight, learned, fighting, understood |
| disney, april, uh, tah, um, phew, god, bo, yep, en, uc, ugh, hannah, su, that’s, oh, ah, montana, i, its, |
| together, mainly, world, majority, many, worlds, maximum, all, both, other, those, theories, correspond, verses, than, well, am, possibly, much, aspects, they’re, call, condition, cause, so, their, saying, mean, no, pieces, as, what, but, too, major, be, if, one, leader, we, easily, have, especially, exactly, list, everyday, not, only, channel, used, make, now, best, area, they, also, because, science, class, our, means, marketing, by, email, areas, are, pursue, applying, that, more, expecially, times, position, rather, though, is, this, us, practice, memory, for, soon, how, possibe, trait, excatly, which, generation, order, it, orders, true, about, being, haven’t, or, association, example, older, long, division, care, either, who, place, were, exact, steps, private, the, process, within |
| chose, going, wants, do, go, depends, can, cant, could, did, don’t, does, let, might, should, will, gonna, would, you’re, get, weralme, wanted, refused, wanna, gets, refuse, take, enough, prefer, able, didn’t, your, you, put, wait, em, else, want |

### Figures

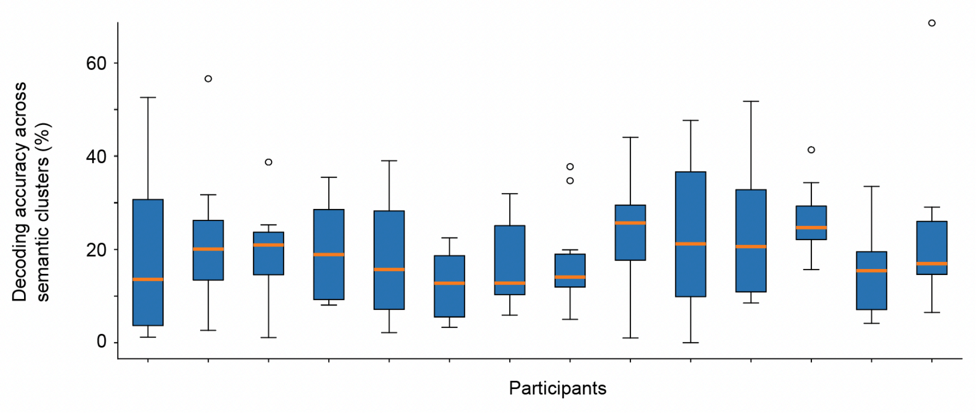


**Figure S1. Decoding accuracy across semantic cluster.** Boxplot showing decoding results for 10 lexical semantic clusters across participants. Boxes represent the distribution of values, including the first and the third quartile across semantic clusters. The whiskers extend from the box label the length of 1.5x the inter-quartile range (IQR) from the box. The orange line indicates the median accuracy across clusters. The empty circle indicates the flier points that passed the range of whiskers.

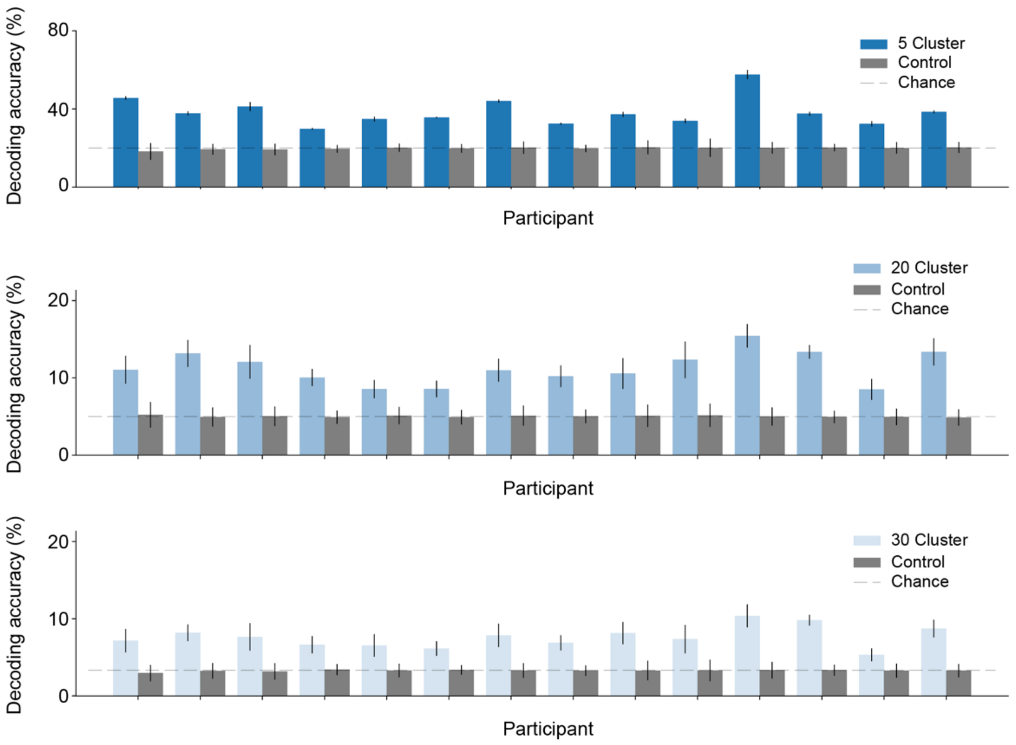


**Figure S2. Lexical decoding at various granularity.** The decoding accuracy from 5 clusters (*top*), 20 clusters (*middle*), and 30 clusters (*bottom*) are compared to random permutation controls (T-test, $p \leqslant 1 \times 10^{-7}$).

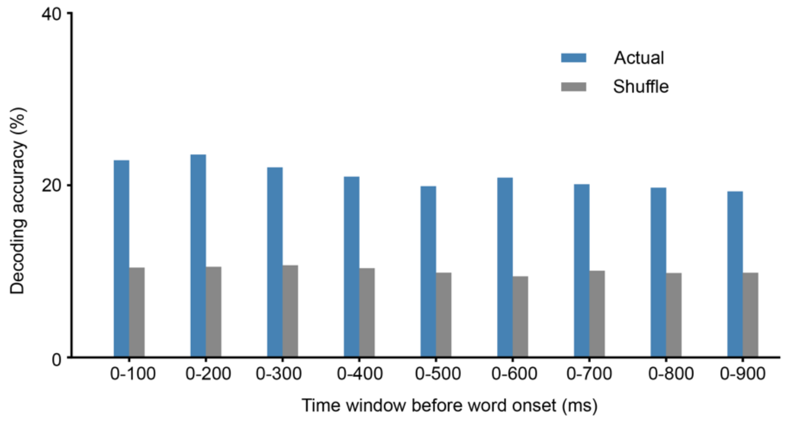


**Figure S3. Decoding accuracy with various time duration.** The decoding accuracy of neural activity were examined across different time windows and consistent performance across these intervals were observed.

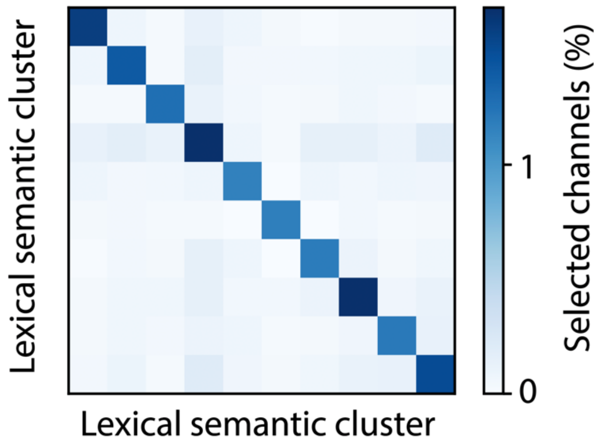


**Figure S4. Channels selecting to one or more lexical semantic clusters.** The bivariant plot shows the percentage of channels with significant response to one or more semantic clusters.

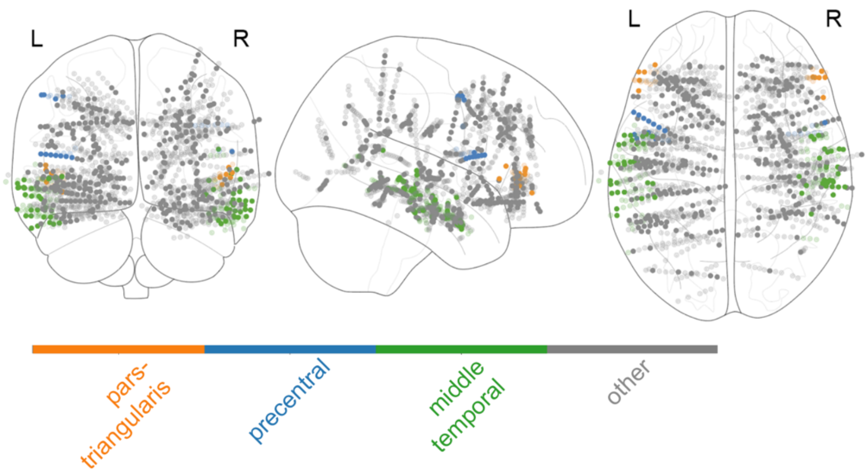


**Figure S5. Illustration of selected channels from pars-triangularis, precentral and middle temporal cortex.** These regions showed significant higher percentage of channels responding to lexical semantic clusters, and the locations of these channels were shown in the figure.

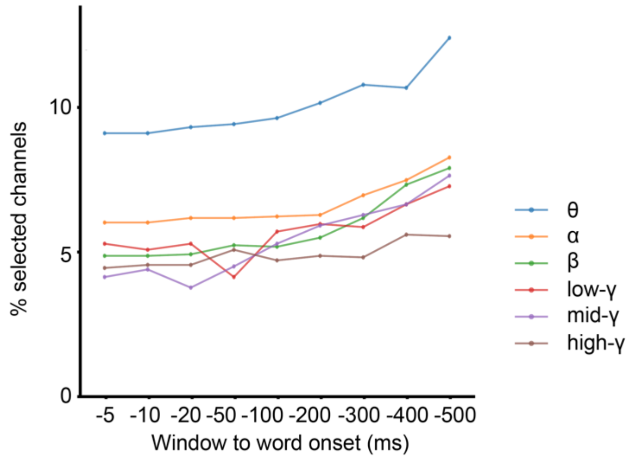


**Figure S6. Percentage of selected channels across time windows and frequency bands.** We examined how different time windows for calculating neural activity affect the percentage of selected channels across various frequency bands. Specifically, these time windows were obtained from the listed time to word onset, which was set as the 0 millisecond. Across all examined time windows, the percentage of channels responding to lexical semantic clusters was consistently highest in the theta frequency band compared to other frequency ranges. This observation suggests that the predominance of low-frequency band responses cannot be attributed to extended time window analysis.
